# Real-time lens-free visualization of strong light scattering by biogenic guanine platelets

**DOI:** 10.1101/2021.04.03.438308

**Authors:** Masakazu Iwasaka

## Abstract

Microscopic observation system without lens has a potential to realize ubiquitous sensing network detecting micro/nano biological hazards to modern society because the lens-free imaging device can provide an extremely compact microscope. In addition to *in slico* micro-mirrors, liquid injectable organic micro-mirrors should be found and utilized for achieving the ideal imaging device for micro/nano objects. This study demonstrates a high contrast lens-free image of the projection from a biogenic guanine platelet floating in water. The fish guanine platelet generated intense and high directional diffraction as well as regular reflection of incident light. The light projection from guanine platelet individually formed an intense platelet-shaped image in real-time on CMOS image sensor arrays. A dynamic projection movie of the guanine platelet, the size of which was approximately 20 ∼ 40 μm × 5 ∼ 10 μm × 100 nm in thickness, was obtained in a small aqueous droplet whose height was less than 2 mm. The developed new lens-free technology using biogenic tiny platelet has an ability to portably visualize movements of the micro/nano objects interacting with the platelet. The compact lens-free inspection can contribute to keep our society in safe.

---

Recently, developments of micro/nano mechanical systems (MEMS/NEMS) brought new technologies such as digital micro-mirror device (DMD) which can be used in various kind of novel display units one of which was head mount display^1^. Dynamic Projection mapping is also a candidate of ubiquitous displaying^2^. DMD is a successful photonic device using a flat mirror with a surface layer providing enough reflectance. However, it should be important to explore different kinds of light controlling materials for the purpose of light projection on micrometer space. Some studies on based on functional materials reported strong light scattering properties of ZnO ^3^ and TiO_2_ ^4^ fine particles. It is hoped to obtain a new efficient photonic device developed by a fusion of novel materials and matured micro manufacturing techniques.

If we switch our viewpoint from artificial techniques to evolutionally achieved techniques in nature, a biomimetic approach to developing and improving a photonic device become possible. Finding a mechanism for light reflection and light scattering utilized by living creatures, which is different from artificial photonics may be beneficial to our society. In the field of ichthyology, there is a long history of research on light reflection by fish body surface.^5^ A tiny platelet made of guanine, which is a DNA base molecule, has a unique optical property as a biogenic micro-mirror plate. ^5-9^

Different from the micro-mirror plate of DMD, biogenic guanine platelets have high transparency for lower incident angles, and at the same time, a guanine platelet show reflectance of more than 20 % for a specific angle of incidence. Furthermore, a stack of guanine platelets with spacings of water exhibited a reflectance of more than 60 %.^9^ It means that the guanine platelets can act as a mirror suitable for bio-microfluidic device.^7^

In addition, recent bio-chip are requiring a new platform of optical microscopy without lens. Because light illumination with lens restricts down-sizing of the bio-chip device, lens-free imaging systems are developed.^10^ Studies on lens-free imaging were carried out on complementary metal oxide semiconductor (CMOS) or charge coupled device (CCD) detectors. By using these imaging devices and computational algorithm, lens-free microendoscope^11^ was developed. Lens-free shadow imaging is one of the promising techniques for ubiquitous medical applications.^12^ Various kinds of image reconstruction methods without lens were proposed on phase retrieval^13^ and new inverse problem algorithms^14,15^. It became possible to apply the lens-free to 3D cell culture^16^.

The present study shows an evidence that the biogenic guanine platelet, which has refractive index of ∼1.83 and thin thickness, exhibits intense projection images on a CMOS image-sensor-array in the absence of lens.

Figure 1 shows the configuration of guanine platelets and lay of incident light on CMOS imager from which lens was detached. The experiments were carried out by using two kinds of commercially obtained CMOS imaging chip. As shown in FIG. 1(a), a drop of aqueous liquid containing guanine platelets from a fish (three species) was directly put on the glass plate covering CMOS imaging chip. An incident light (from LED or laser) was supplied from side. Figure 1(b) illustrates the condition of light propagation inside the droplet forming a dome due to surface tension. The ray of incident light was adjusted not to directly illuminate the pixels of CMOS chip. Figure 1(c) shows an image of fish guanine platelet. Typical size of main face of the biogenic guanine platelet is 20 ∼ 40 μm × 5 ∼ 10 μm. Thickness of the platelet was 100 nm ∼ 160 nm. This thin platelet floated in water for an hour and made it possible to control the propagation of light.

**FIG. 1.**
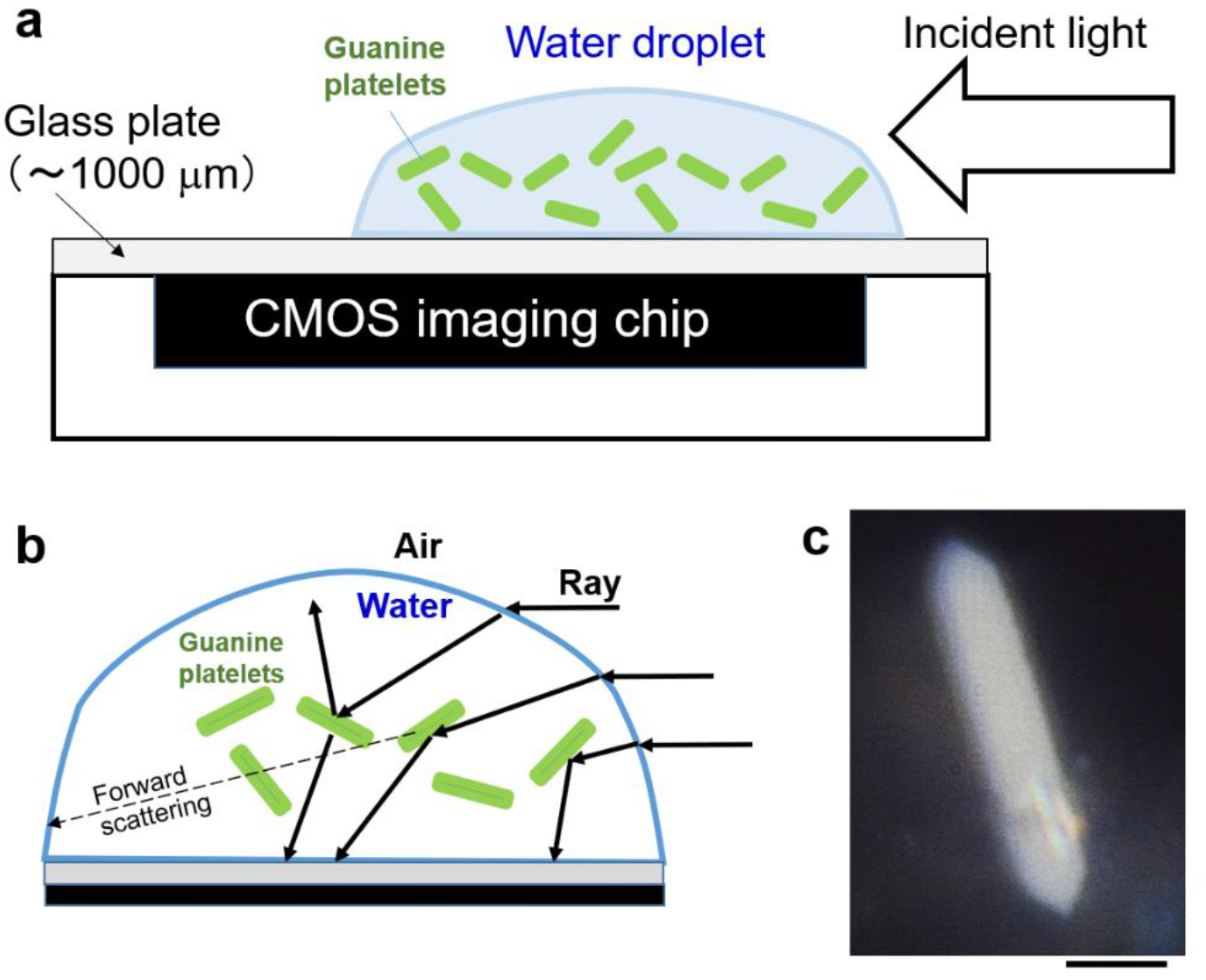
Observation system for light scattering by guanine platelets in water utilizing CMOS image sensor chip without lens. (a) Experimental diagram of the liquid sample on CMOS imaging chip with glass plate under exposure to incident light. (b) Illustration of lays which is reflected and diffracted by a guanine platelet floating in water. (c) Optical microscopic image of a biogenic guanine platelet floating in water. Scale bar is 10 μm.

Figure 2 shows the direct projection of the floating platelet’s light scattering on the CMOS chip in the absence of lens. The light source was white LED. In the case of Fig. 2(a), the liquid droplet was covering whole area of the cover glass plate of chip. The incident light came into the dome via air-water boundary, and reached a guanine platelet floating in water. Figure 2(b) and (c) are the cases when boundary of air and liquid existed upon the imager plate. The setting was carried out to make a distinction of distribution of projected images between inside and outside the dome. Most of the individually projected images appeared right beneath the dome. In the left side of the dome, strong light reflection by water-air boundary occurred. There were some images projected on outside of the dome (left side of FIG. 2(c)).

**Figure 2.**
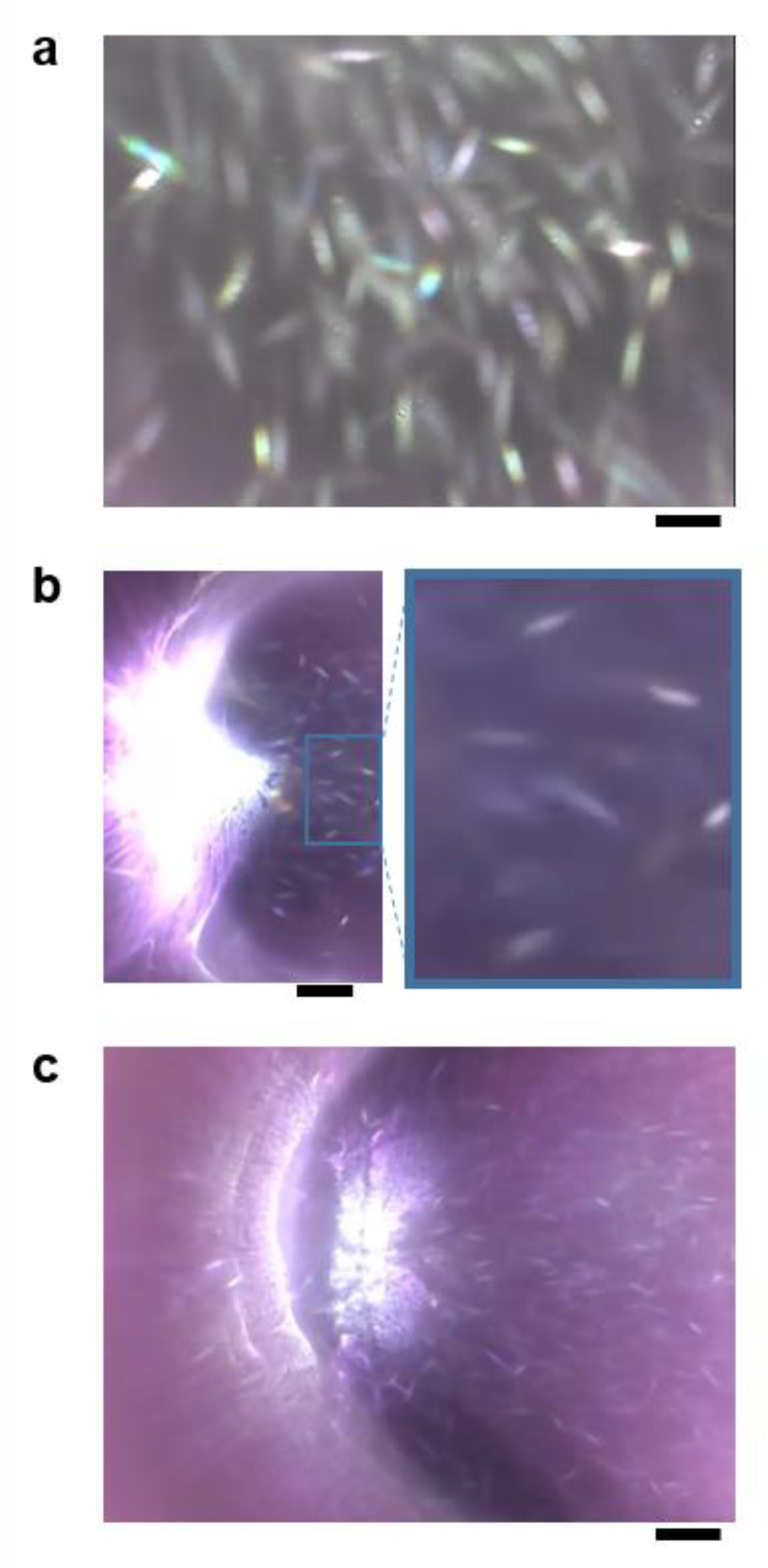
Lens-free imaging of individually projected pattern of light which was scattered by a guanine platelet floating in water. (a) Image of projection when water droplet covering whole area of CMOS chip. Guanine platelet is of goldfish, *Carassius auratus*. (b) Water droplet covered half area. Projection under the droplet is expanded in right panel. (c) Individually projected light scattering from a platelet also appeared in the outside of water droplet covering area (left hand side of the water-air boundary). Guanine platelet of Japanese anchovy, *Engraulis japonicus* was used in b and c. Scale bar is 400 μm.

The biogenic guanine platelets in water exhibited a very curious phenomenon, frequently moving projection images, as shown in a movie of Supplementary Materials. Previous studies suggested that the guanine platelets caused light flickering due to rotational motion arising from thermal agitation in water. Thickness of the liquid dome was less than 2 mm, and thickness of glass plate covering CMOS pixels was approximately 1 mm. Images shown in FIG. 2 indicated that the size of individual platelet image was 200 μm∼400 μm. The real size of the platelet was 20 μm∼40 μm in maximum length. Roughly estimating, the individual projection 10 times expanded in a propagation distance of 2 mm. In this condition, a slight tilting of a platelet can cause a long-distance movement of the projected image on imaging plane.

Next, dependence of light wavelength on the formation of projection was investigated by using three laser lights, in blue, green and red. Figure 3 shows the lens-free imaging of the laser light scattering patterns. In the comparison of projected patterns, green and blue light scattering exhibited a clear projection on the CMOS imager plane, while obscure projections were obtained under a red laser light. Particularly, compared to blue light and red light, the green laser light provided stronger and larger patterns of projection.

**Figure 3.**
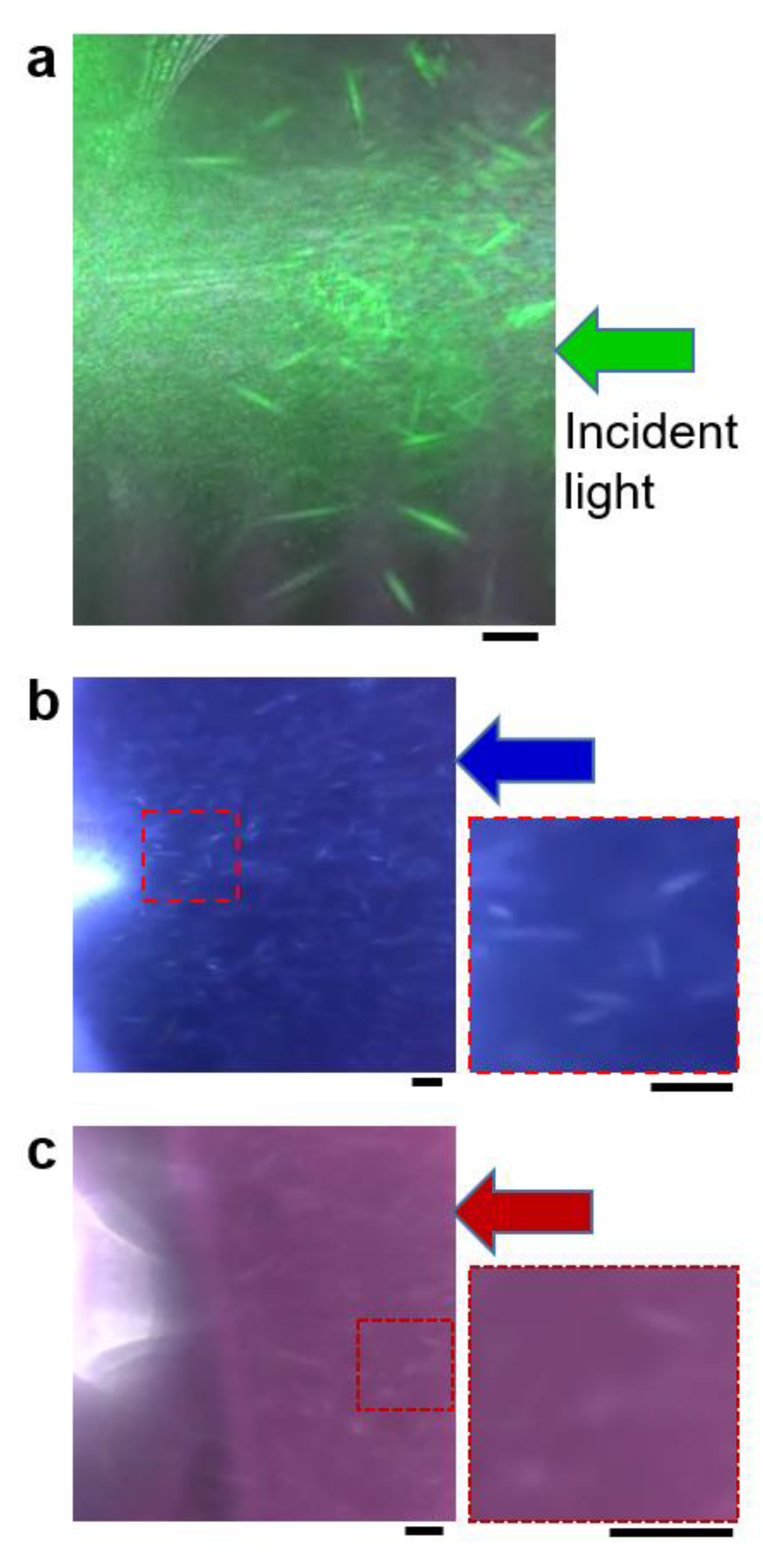
Lens-free imaging of green, blue and red laser light scattering. (a)Under exposure to green laser light at 530 nm. (b) Under exposure to blue laser light at 480 nm. (c) Under exposure to red laser light at 630 nm. Right panel of (b) and (c) shows a magnification of the photograph. Guanine platelet of yellowback seabream, *Dentex tumifrons* was utilized. Scale bar is 200 μm.

In order to understand the mechanism of scattering pattern formation which was directly projected on CMOS imaging plate without lens, a numerical simulation employing FDTD (finite-difference time-domain) method’s algorism was carried out for analyzing light scattering by a guanine platelet, as shown in FIG. 4.

**Figure 4.**
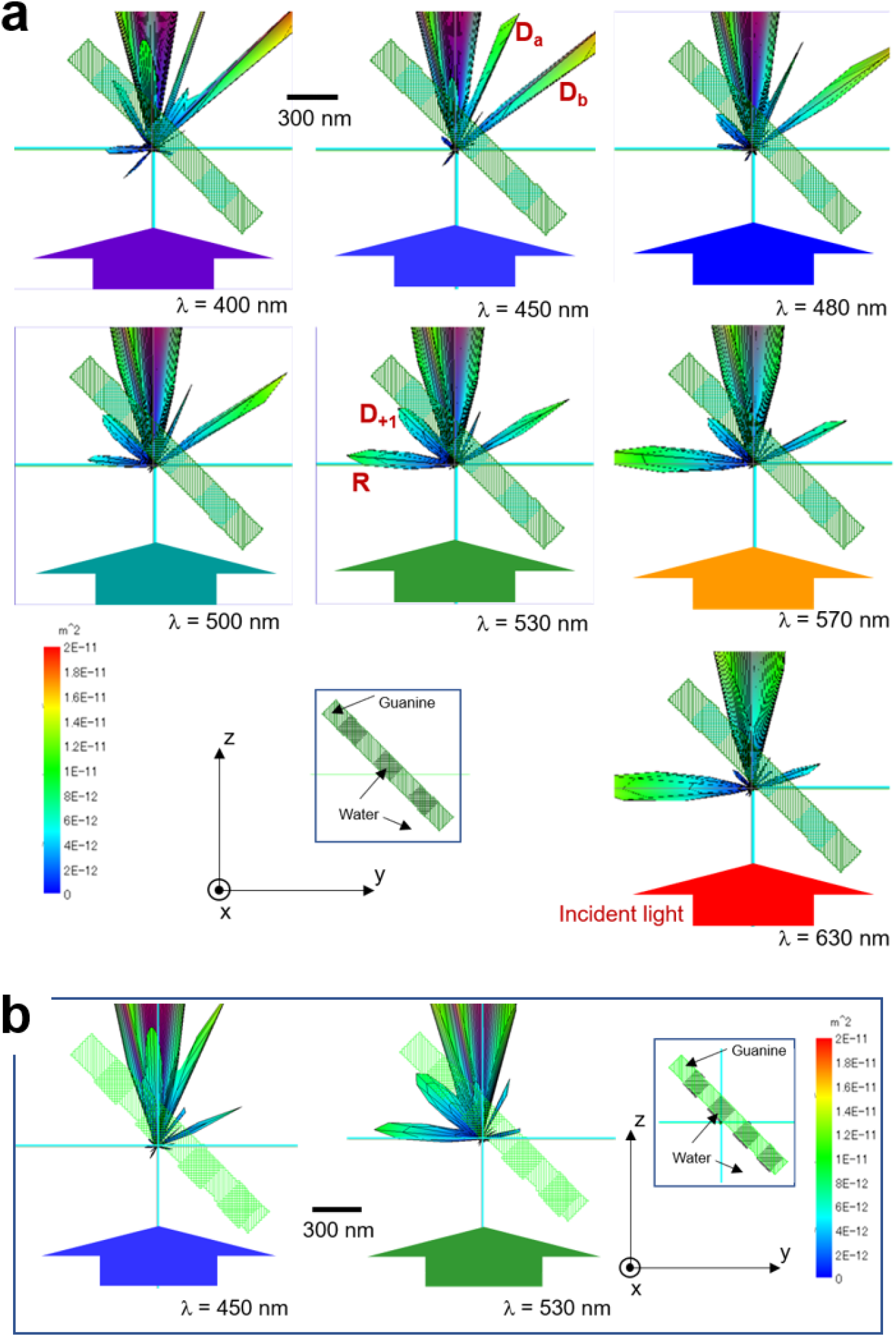
Numerical (FDTD) simulation of light scattering by a guanine platelet having nano-hole grating. The platelet is floating in water and tilts by 45° versus the incident light direction. The incident light is *s*-polarized when incident plane is set to be y-z plane. Size of the platelet is 1381 nm× 1395 nm× 160 nm. The nano-holes in platelet are filled with water and the platelet (a) Guanine platelet with grating period of about 400 nm. (b) Guanine platelet with grating period of about 300 nm. Detailed arrangement of the nanoholes in platelet is shown in supplementary materials. The vertical color indicator shows area of cross section (m^2^). The maximum value of the cross section is fixed at 2 × 10^−11^ (m^2^) in all graphs.

In the previous study on internal structure of fish guanine platelet, it was found that grating structures in several hundreds of nanometers existed in the platelet.^17^ The simulation was conducted by following this kind of structure. The model of the platelet was consisted of a guanine (refractive index = 1.83) and nano-hole grating. Water (refractive index = 1.33) surrounded the platelet and existed inside the nanoholes. By changing the distance of nano-holes, i.e. period of grating and tilting angle versus incident light, scattering patterns in the guanine platelet model were evaluated (Figure 4 and Supplementary Materials).

In FIG. 4(a) where the grating distance was approximately 400 nm, the incident light propagating to +*z* axis has an incident angle of 45° versus the platelet in water. The scattering patterns for lights at seven wavelengths had a regular reflection (denoted by R) except for the light at 450 nm. On the contrary, two intense diffractions, D_a_ and D_b_ occurred at *λ* = 450 nm. Intensities of the diffraction D_b_ were greater than the regular reflection R at *λ* = 400 nm ∼ 530 nm. In addition, a diffraction (denoted by D_+1_) directing in parallel to the platelet’s maximum plane became distinct at *λ* = 450 nm ∼ 570 nm. Diffractions D_+1_ and D_b_ were also observed when the refractive index of platelet was changed to 1.5 and the thickness was 210 nm, although their intensities were smaller (*see* Supplementary Materials). So, it was conjectured that diffractions D_b_ originated from the nano-hole grating in guanine platelet.

The switching of diffraction and reflection was remarkable when comparing the pattern at *λ* = 400 nm and 530 nm. This tendency remained even when the grating distance became shorter (∼ 300 nm) and the tilting angle of platelet varied, as shown in FIG. 4(b) (and Supplementary Materials).

In comparison with the biogenic guanine platelets, micro/nano particles of hexagonal boron nitride, barium sulfate and titanium oxide did not provide any projected pattern on the CMOS imager plate without lens under white LED light or laser light. Higher reflectance was not efficient for realizing the lens-free projection on CMOS chip. Light scattering directionality of the particle and reducing random scattering were important for obtaining the high contrast visualization of the projection without lens. Thin thickness of the biogenic guanine platelets and nano-holes filed with water should have a role to project both reflection and diffraction separately from the forward scattering which was in parallel to incident light. Previous works of lens-free system were setting the light source unit over image chip that made the apparatus thicker. We can expect to reduce the distance of light source and image sensor array by employing the guanine platelets in aqueous liquid existing near the target sample on chip. In conclusion, the guanine platelets derived from fish had a remarkable ability to form a high contrast image of projection on imaging plane of CMOS sensor arrays.

## AUTHORS’ CONTRIBUTIONS

M.I. performed the experimental design, experiments, measurements and analyses. All part of manuscript and illustrations were prepared by M. I.

## ACKNOWLEDGMENTS

This work was supported by JST-CREST “Advanced core technology for creation and practical utilization of innovative properties and functions based upon optics and photonics (Grant number: JPMJCR16N1).”

## DATA AVAILABILITY

The data that support the findings of this study are available from the corresponding author upon reasonable request.

## SUPPLEMENTARY MATERIAL

### 1. Real-time movies of guanine platelet projecting light on CMOS image sensor array without lens

#### Descriptions of experimental materials

- CMOS image sensor-A: CMOS interface module CM405-OA (Asahi Electronics, Osaka, Japan) Presented data: Figure 2a
- CMOS image sensor-B: USB camera module ELP-SUSB1080PO1-L36 (Ailipu Technology Co., Ltd, Shenzhen, Guangdong, China) Presented data: Figures 2b-2c, Figures 3a-3c

#### Additional information of the lens-free observation

- The lens provided to the above mentioned two CMOS module was detached, and glass plate covering image sensor was opened to the air, as shown in FIG.1.
- The movie of the lens-free image was recorded in PC to which COMS module connected

#### Biological sample

- Biogenic guanine platelets were obtained from scale or skin of goldfish, *Carassius auratus*, Japanese anchovy, *Engraulis japonicus* and yellowback seabream, *Dentex tumifrons* (obtained from food market). The experimental methods of the treatment of fish were approved by Hiroshima University Animal Care and Use Committee (approval number: F16–2 and F20-4, Hiroshima University).
- The separated guanine platelets were suspended in distilled water, and diluted to be a density of approximately 10^3^ per ml when used for the lens-free observation. 40 ml of the suspension containing platelets was put on the glass surface covering the CMOS image sensor via a micro-pipette
- The surface of glass had a hydrophobic property without any additional hydrophobic treatment, and a dome of aqueous liquid was obtained.

**Figure2aMovie: Real-time lens-free imaging of projected white light from guanin platelets** (guanine platelet of goldfish).

This movie provided the images for Figures 2a. White light illumination is supplied from right side of the picture.

**Figure2bMovie: Real-time lens-free imaging of projected white light from guanin platelets** (guanine platelet of Japanese anchovy). Boundary of water and air is set in the image sensor area in order to discriminate the projection existing inside the dome.

This movie provided the images for Figures 2b. White light illumination is supplied from right side of the picture.

**Figure2cMovie: Real-time lens-free imaging of projected white light from guanin platelets** (guanine platelet of Japanese anchovy). Boundary of water and air is set in the center of image sensor area in order to observe the individually projected image of platelet existing outside the dome. This movie provided the images for Figures 2c. White light illumination is supplied from right side of the picture.

**Figure3aMovie: Real-time lens-free imaging of projected green laser light from guanin platelets** (guanine platelet of yellowback seabream). This movie provided the images for Figures 3a. Green laser light is supplied from right side of the picture.

**Figure3bMovie: Real-time lens-free imaging of projected blue laser light from guanin platelets** (guanine platelet of yellowback seabream). This movie provided the images for Figures 3b. Blue laser light is supplied from right side of the picture.

**Figure3cMovie: Real-time lens-free imaging of projected red laser light from guanin platelets** (guanine platelet of yellowback seabream). This movie provided the images for Figures 3c. Red laser light is supplied from right side of the picture.

The original movies were in AVI format, but the files were converted to MPEG-4.

### 2. Supplementary data and model of FDTD simulation

FDTD (Finite-difference time-domain) method simulation was carried out to obtain light scattering pattern in a platelet floating in water. Electromagnetic fields of a pulsed light propagating to +z. A commercial product of electromagnetic fields solution software, Poynting for Optics V03L10R121 (Fujitsu co. ltd., Tokyo, Japan) was utilized. After calculating near field electromagnetic fields of the light propagated around the platelet,

This light source had a rectangular volume and propagated inside a rectangular box filled with water (refractive index n = 1.33). The platelet (n = 1.83 for guanine) had a thickness of 160 nm and was set inside the rectangular box of water. The detailed condition of the incident light was explained in the main text. Optical wavelengths of the pulsed light for monitoring were set at 400 nm, 450 nm, 480 nm, 500 nm, 530 nm, 570 nm, and 630 nm.

FigS4-1 shows the model for FDTD simulation shown in Figure 4b. FigS4-2 ∼ FigS4-4 show spherical patterns of light scattering at three angles of incident light (60°, 30°, 45°). FigS4-5 shows spherical patterns of light scattering by a platelet with refractive index of 1.5.

**Fig S4-1.**
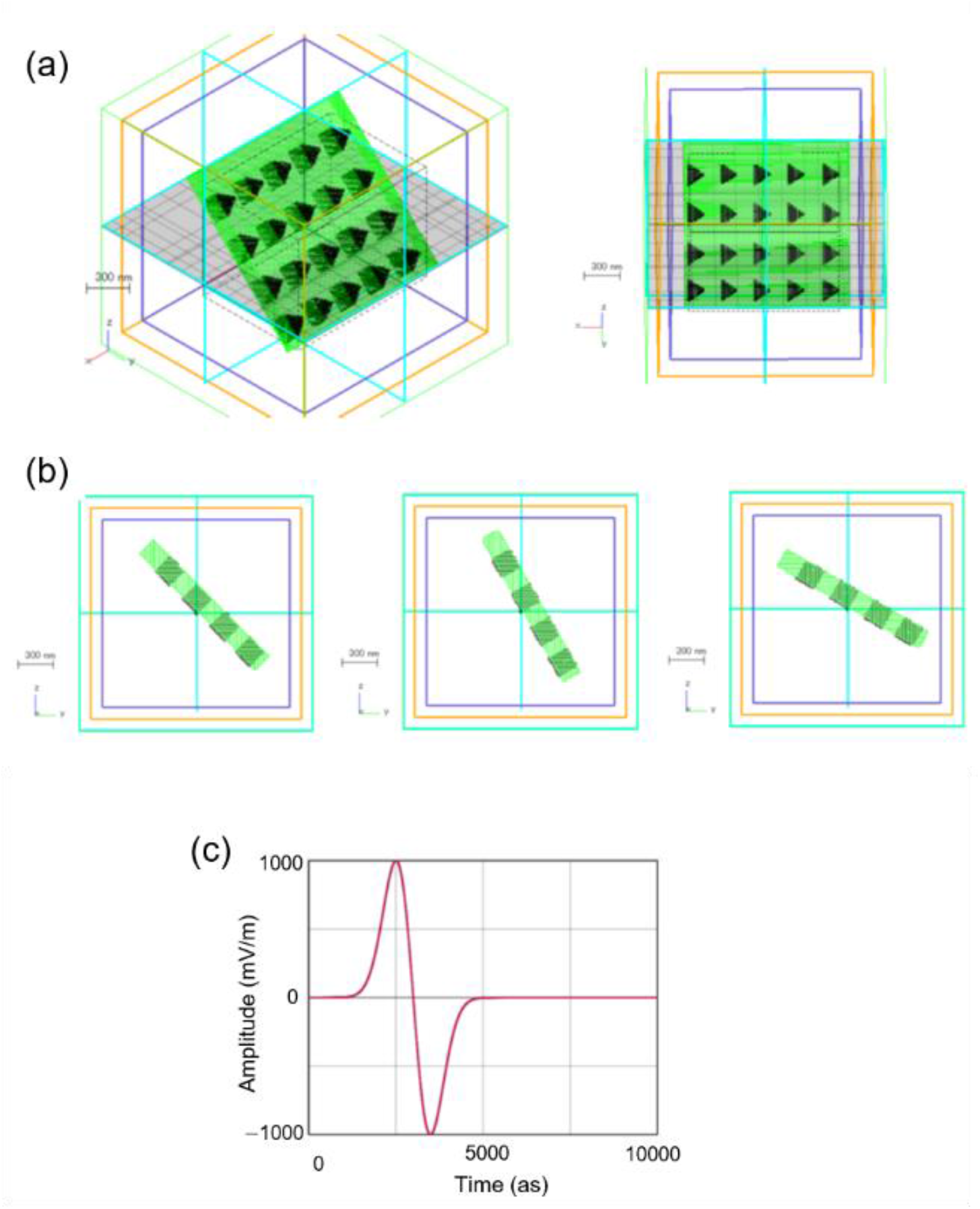
Model of guanine platelet with nano-holes utilized for the FDTD simulation shown in Figure 4b. (a) Overview of the model. (b) Side view at three angles of tilt (three angles of incident light (60°, 30°, 45°)). (c) Waveform of incident light (pulsed light having components of optical wavelengths for analyzing). The same light source was utilized for the simulation shown in Figure 4a and FigS4-2 ∼ FigS4-5.

**Fig S4-2.**
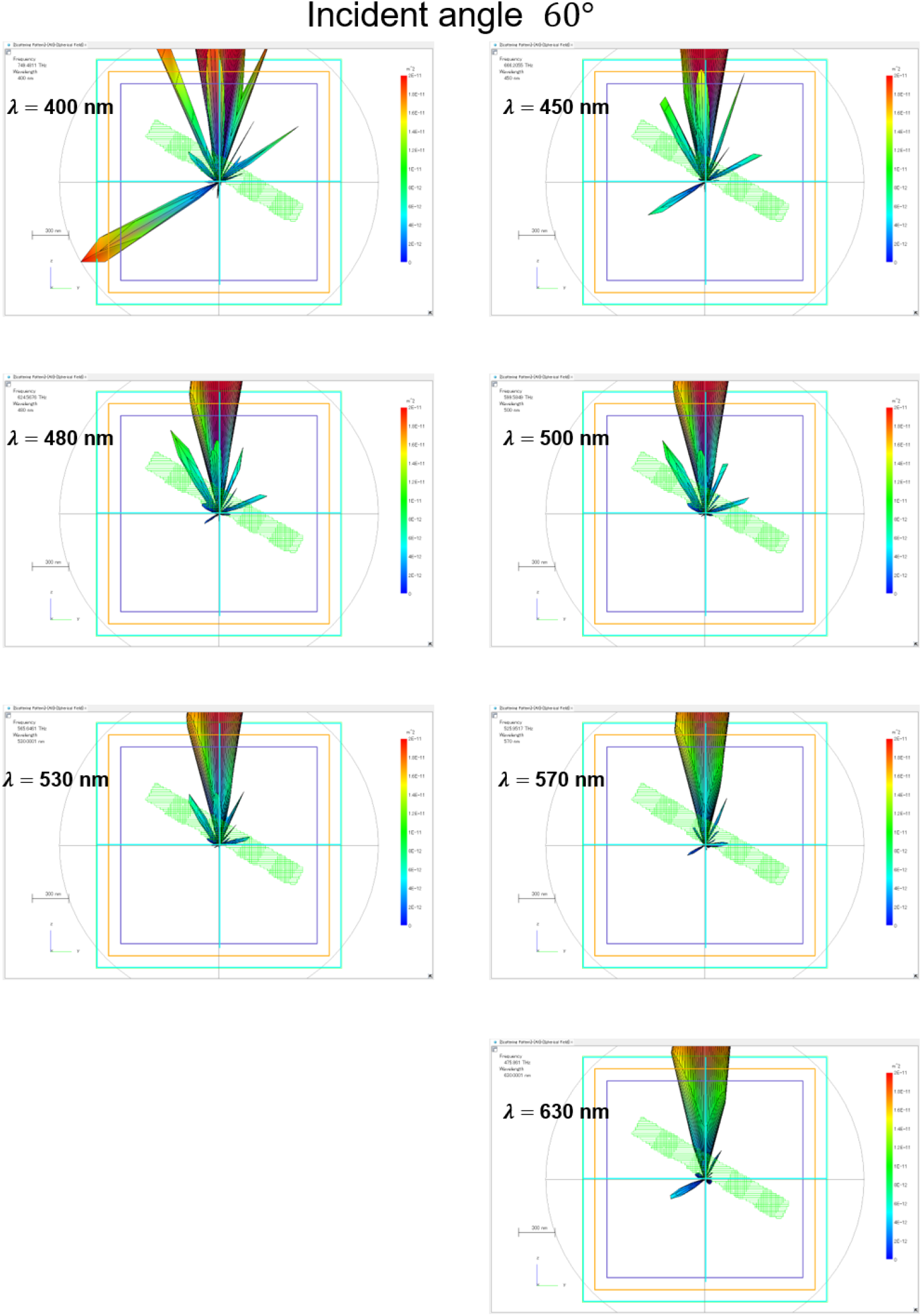
FDTD simulation of light scattering by a guanine platelet model shown in FigS4-1a when the platelet tilts by 60° versus the incident light direction.

**Fig S4-3.**
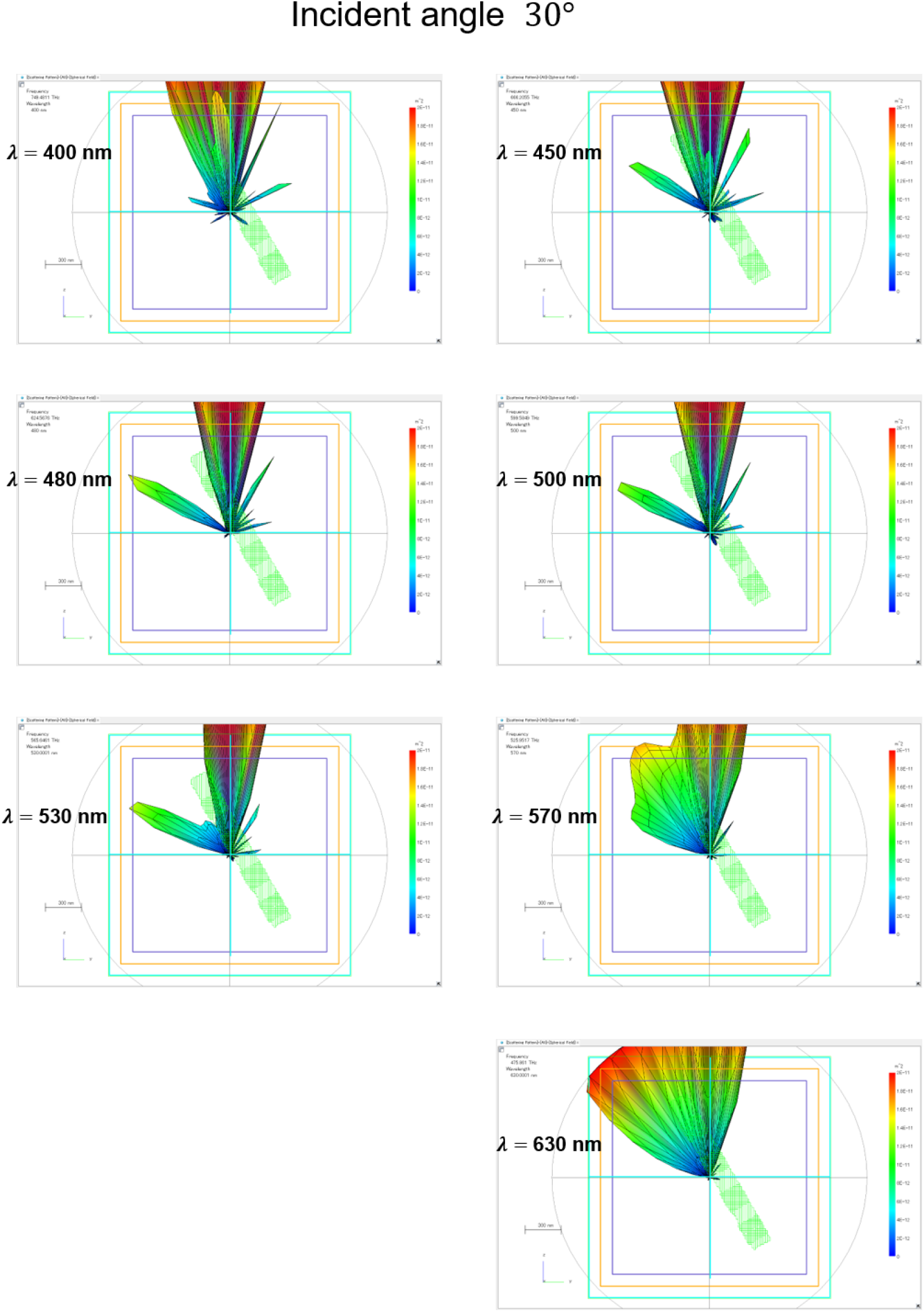
FDTD simulation of light scattering by a guanine platelet model shown in FigS4-1a when the platelet tilts by 30° versus the incident light direction.

**Fig S4-4.**
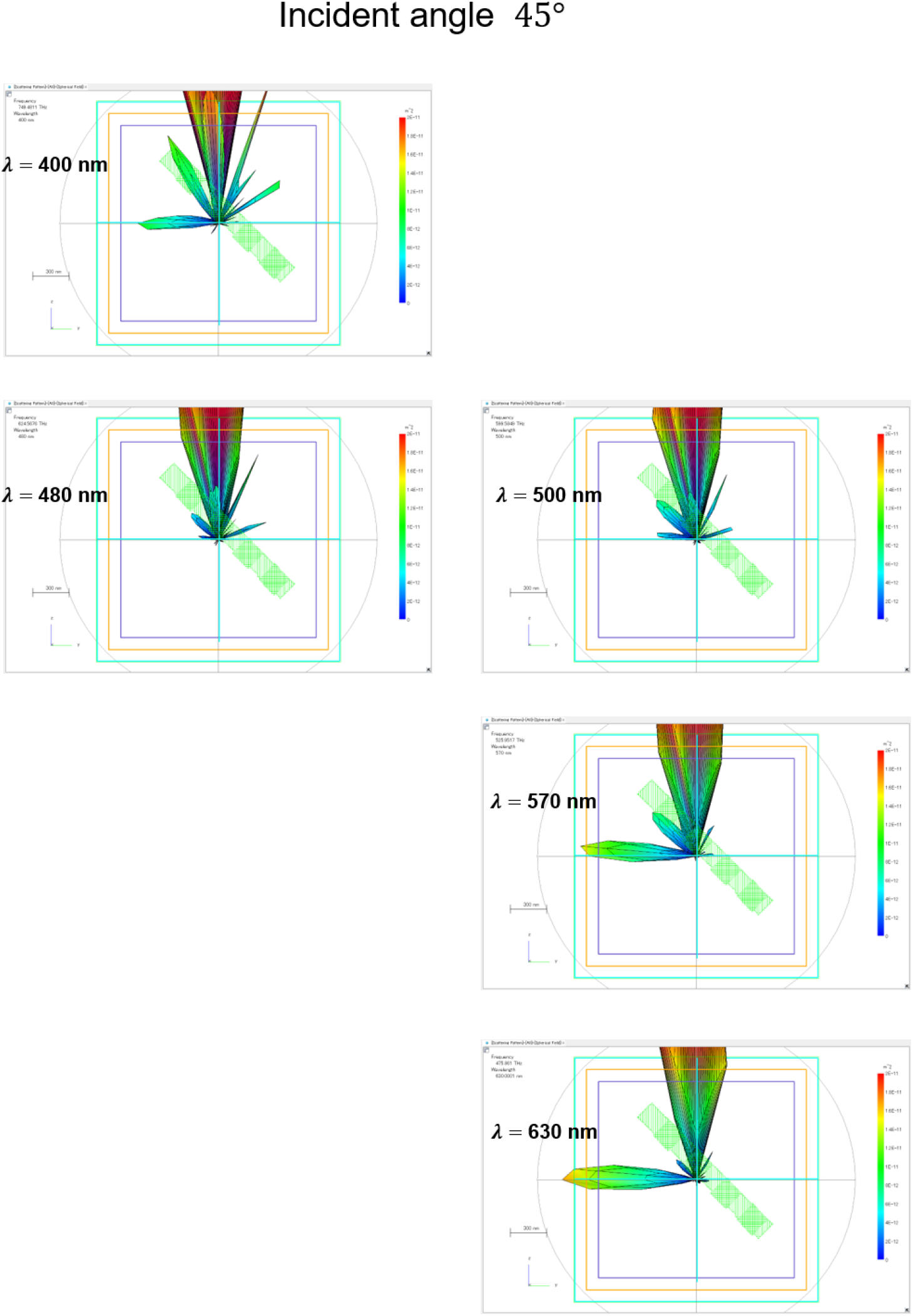
Complemental data for Figure 4. The platelet tilts by 45° versus the incident light direction. Spherical plots of light scattering by platelet at 400 nm, 480 nm, 500 nm, 570 nm and 630 nm are shown. The data at 450 nm and 530 nm are shown in Figure 4 of the main text.

**Fig S4-5.**
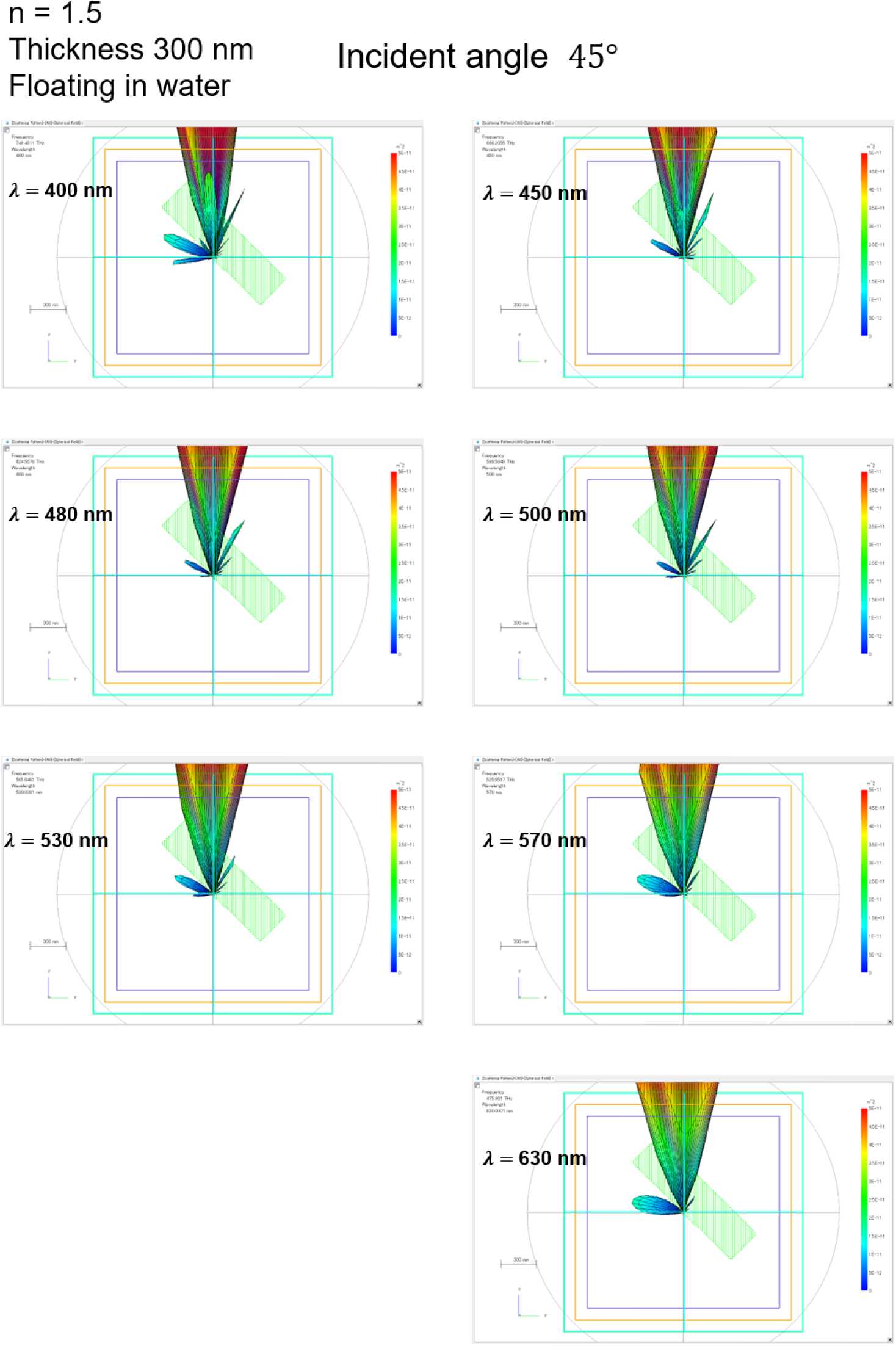
FDTD simulation of light scattering by a platelet with refractive index of 1.5 which were tilting by 30° versus the incident light direction. Thickness of the platelet: 210 nm

